# Chronic inflammatory pain drives alcohol drinking in a sex-dependent manner

**DOI:** 10.1101/379362

**Authors:** Waylin Yu, Lara S. Hwa, Viren H. Makhijani, Joyce Besheer, Thomas L. Kash

## Abstract

Sex differences in chronic pain and alcohol abuse are not well understood. The development of rodent models is imperative for investigating the underlying changes behind these pathological states. However, past attempts have failed to produce drinking outcomes similar to those reported in humans. In the present study, we investigated whether hind paw treatment with the inflammatory agent Complete Freund’s Adjuvant (CFA) could generate hyperalgesia and alter alcohol consumption in male and female C57BL/6J mice. CFA treatment led to greater nociceptive sensitivity for both sexes in the Hargreaves test, and increased alcohol drinking for males in a continuous access two-bottle choice (CA2BC) paradigm. Regardless of treatment, female mice exhibited greater alcohol drinking than males. Following a 2-hour terminal drinking session, CFA treatment failed to produce changes in alcohol drinking, blood ethanol concentration (BEC), and plasma corticosterone (CORT) for both sexes. 2-hr alcohol consumption and CORT was higher in females than males, irrespective of CFA treatment. Taken together, these findings have established that male mice are more susceptible to escalations in alcohol drinking when undergoing pain, despite higher levels of total alcohol drinking and CORT in females. Furthermore, the exposure of CFA-treated C57BL/6J mice to the CA2BC drinking paradigm has proven to be a useful model for studying the relationship between chronic pain and alcohol abuse. Future applications of the CFA/CA2BC model should incorporate manipulations of stress signaling and other related biological systems to improve our mechanistic understanding of pain and alcohol interactions.

## Introduction

Alcohol provides an accessible means of self-medication for pain suffering populations, with approximately 25% of chronic pain patients consuming alcohol for symptom relief (Riley and King, 2009). Although alcohol can act as a transient pharmacological intervention for pain, excessive drinking can lead to increased risk for addiction and related heath complications (Koob and Volkow, 2010). Chronic exposure to alcohol is especially detrimental for pain, with marked increases in sensitivity after drug cessation (Boissoneault et al., 2018; Dhir et al., 2005; Dina et al., 2000, 2007; Dodds et al., 1945; Edwards et al., 2012; Fu et al., 2015; Gatch and Lal, 1999; Jochum et al., 2010; Malec et al., 1987; Riley and King, 2009; Roltsch et al., 2017; Shumilla et al., 2005; Smith et al., 2015; Wolff et al., 1942). Clinical observations have suggested important differences in how men and women consume alcohol and experience the detrimental consequences of alcohol abuse when suffering from pathological pain. Among chronic pain patients, alcohol use is more prevalent in males (Brennan et al., 2011; Egli et al., 2012; Riley and King, 2009; Wilsnack et al., 2009). Females that habitually consume alcohol, however, are more susceptible to pathological pain (Boissoneault et al., 2018). To date, preclinical investigations have failed to model similar sex-specific patterns in drinking and pain, thus providing a major hurdle to studying the mechanisms of pain-related alcohol consumption (Smith et al., 2015; Yezierski and Hansson, 2018).

To address this deficit in preclinical models, the present study examines alcohol consumption in male and female C57BL/6J mice treated with Complete Freund’s Adjuvant (CFA), an antigen emulsion containing heat-dried *Mycobacterium tuberculosis*. CFA was selected to model chronic inflammatory pain, since it closely mimics the time course of persistent injury (Hargreaves et al., 1988; Larson et al., 1986; Ren and Dubner, 1999). Furthermore, postoperative pain, arthritis, and related inflammatory states have exhibited a close relationship with alcohol abuse (King et al., 2012). Therefore, we hypothesized that CFA-treated mice would show increased alcohol consumption compared to uninjured mice. To test this, we exposed saline- and CFA-treated mice of both sexes to a voluntary drinking paradigm (i.e. CA2BC with 20% ethanol [EtOH]) and measured alcohol consumption across a three-week timespan. Thermal nociceptive thresholds were assessed to ensure that CFA treatment altered pain sensitivity. Blood EtOH concentration (BEC) and plasma corticosterone (CORT) levels were analyzed to assess pharmacologically relevant drinking and alcohol-related stress signaling, two possible contributors to sex differences in alcohol and pain interactions (Edwards et al., 2012; Egli et al., 2012; Fu and Neugebauer, 2008; Heilig and Koob, 2007; Koob, 2013). Taken together, these experiments aim to successfully model sex differences in pain-related alcohol drinking for mice, with the hope that it can be applied towards mechanistic investigations of pain and alcohol interactions in the future.

## Materials and Methods

### Animals

A total of 32 male and female C57BL/6J mice (Jackson Laboratories, Bar Harbor, ME) arrived at 6-weeks of age and were group-housed for one week to habituate to vivarium conditions. Following habituation, mice were individually housed in ventilated Plexiglas cages and given three days of acclimation in a 12-hr reverse light/dark cycle (12am-12pm) prior to the start of the experiment. Subjects were provided *ad libitum* access to water and Isopro RMH 3000 chow (Prolab, St. Louis, MO) for the duration of the experiment. All procedures were approved by the University of North Carolina at Chapel Hill Institutional Animal Care and Use Committee and were in accordance with the NIH Guide for Care and Use of Laboratory Animals.

### Continuous access 2-bottle choice drinking

Male (N = 16) and female (N = 16) mice were given continuous access to 20% EtOH (w/v) and tap water for three weeks. EtOH solutions were prepared with tap water and 95% EtOH (Pharmaco-AAPER, Brookfield, CT), and delivered via sipper tubes made from 50 mL plastic tubes (Nalgene), rubber stoppers (Fisher Scientific, Agawam, MA), and sipper tubes (Ancare Corp., Bellmore, NY). Prior to EtOH exposure, subjects were given three days to adapt to drinking with sipper tubes. Individually housed mice were then given access to two sipper tubes per cage: one containing 20% EtOH and the other containing tap water. For the span of three weeks, EtOH and water bottles were weighed daily three hours into the dark cycle. Bottle placement (left vs. right) was alternated weekly to control for an inherent or developed side preference. To control for non-consumption-related fluid loss, a dummy cage containing identical sipper tubes was used to subtract weekly dripped fluid values from subject fluid consumption.

### Chronic inflammatory pain

Prior to EtOH exposure, subjects were given a 50 μl subcutaneous injection of saline or Complete Freund’s Adjuvant (CFA; Sigma, St. Louis, MO) in the plantar surface of the right hind paw (n = 8). Drinking experiments started two days after paw injections, around the time that CFA exhibits maximum inflammatory hyperalgesia (*unpublished*). To verify that changes in drinking behavior resulted from differences in pain sensitivity, thermal nociceptive thresholds of saline- and CFA-treated mice were assessed one day after the last 24-hr EtOH exposure using the Hargreaves test. Mice were placed in a plexiglas chamber elevated on a glass plate (IITC Life Science Inc., Woodland Hills, CA) and habituated to the behavioral apparatus for a minimum of 30 minutes. The mid-plantar surface of saline/CFA-treated and untreated hind paws was then exposed to three heat trials each and assessed for paw withdrawal latencies (PWL). The beam intensity was set to produce basal PWLs of approximately 5-7 seconds, with a maximal value of 20 seconds to prevent excessive tissue damage. All testing was conducted with investigators blinded to the experimental conditions.

### Blood ethanol and corticosterone concentrations

Following pain sensitivity testing, mice were given four days to recover and re-establish drinking behavior in preparation for blood EtOH and corticosterone measurements. Mice were sacrificed 2 hours after sipper tubes were presented, with trunk blood immediately collected in centrifuge tubes following decapitation. Blood samples were centrifuged at 4°C for 10 min at 3000 RPM, and the plasma was separated for storage at −80°C until analysis. To measure blood EtOH concentration (BEC; mg/dl), 5 μl plasma samples were administered through a Model AM1 Alcohol Analyser (Analox Instruments Ltd., Lunenburg, MA). To measure plasma corticosterone (CORT; ng/ml), 5 μl plasma samples were processed with a commercially available colorimetric ELISA kit (Arbor Assays; Ann Arbor, MI), according to the manufacturer’s instructions, with all samples run in duplicate.

### Statistical analysis

All statistical analyses were performed using Prism 6 (GraphPad Software Inc., La Jolla, CA). Analysis of drinking across sessions for saline- and CFA-treated mice (i.e. Treatment x Session for EtOH Intake, EtOH Preference Ratio, and Total Fluid Intake) was performed with a repeated measures analyses of variance (RM-ANOVA). Drinking values for each mouse were averaged across the three-week drinking period to compare Treatment x Sex with additional two-way ANOVAs. Differences in PWL were determined with a two-way ANOVA comparing Treatment x Paw. Post hoc analyses with Sidak or Tukey adjustment were performed following significant main group effects. Data are reported as mean plus or minus the standard error of the mean (M ± SEM). Correlational analyses were conducted using linear regressions to assess the predictive relationship of EtOH intake and thermal nociceptive sensitivity, BEC, and CORT levels, with slopes and intercepts assessed for deviation from zero to determine statistical significance.

## Results

### CFA-treated mice exhibit greater alcohol drinking in a sex-dependent manner

#### EtOH Intake

Male mice treated with CFA consumed a significantly greater amount of EtOH per day (g/kg/24hr) than saline-treated controls, while females did not exhibit any consumption differences across treatments (Fig. 1). A RM-ANOVA (Treatment x Session) revealed a main effect for Treatment [*F*(14, 266) = 4.768, p = 0.0465] and Session [*F*(19, 266) = 1.656, p = 0.0437] in males (Fig. 1A), and a main effect for Session [*F*(19, 266) = 9.470, p < 0.0001] in females (Fig. 1B), but no significant interaction for either sex. Sidak post-hoc analysis revealed no significant difference in EtOH intake between saline- and CFA-treated mice during individual drinking sessions of males or females. Mean drinking values for individual subjects were averaged across three weeks in their respective treatment and sex groupings. A two-way ANOVA (Treatment x Sex) revealed a main effect for Treatment [*F*(1, 76) = 26.04, p < 0.0001] and Sex [*F*(1, 76) = 179.6, p < 0.0001], and a significant interaction between Treatment and Sex [*F*(1, 76) = 16.45, p = 0.0001], where female mice exhibit greater EtOH intake than males, regardless of paw treatment (Fig. 1C). Tukey post-hoc analysis revealed a significant difference in EtOH intake between saline- and CFA-treated mice for males, but not females (Fig. 1C).

**Figure 1.**
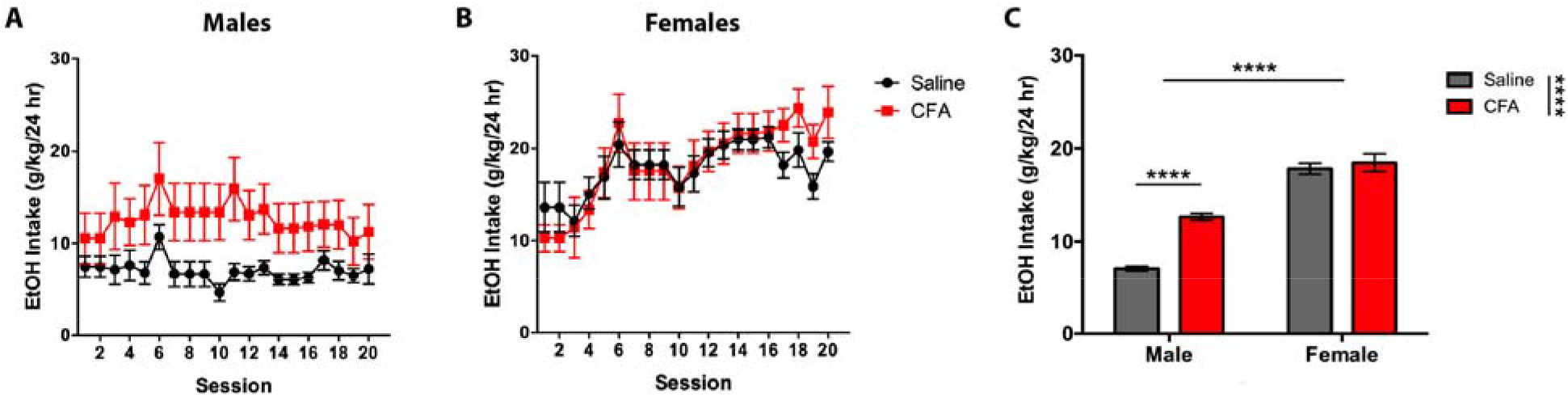
Daily alcohol intake (g/kg/24 hr) of saline- or CFA-treated C57BL/6J mice over a 3-week continuous access regimen. Pain-related consumption of 20% EtOH (w/v) was assessed for (A) males (n = 8) and (B) females (n = 8) in 20 consecutive sessions. (C) Averages of daily intake were compared between saline- and CFA-treated males and females. Data are mean alcohol intake ± SEM. **** p < 0.0001 difference between groups.

#### EtOH Preference Ratio

Preference for EtOH exhibited similar results, with CFA-treated male mice showing a significantly greater EtOH preference ratio than their saline-treated counterparts, while females did not exhibit a preference difference across treatments (Fig. 2). A RM-ANOVA (Treatment x Session) revealed a main effect for Treatment [*F*(1, 14) = 6.777, p = 0.0208] in males (Fig. 2A) and Session [*F*(19, 266) = 5.297, p < 0.0001] in females (Fig. 2B), but no significant interaction for either sex. Sidak post-hoc analysis showed no significant difference in EtOH preference ratio between saline- and CFA-treated mice during individual drinking sessions of males or females. A two-way ANOVA (Treatment x Sex) revealed a main effect for Treatment [*F*(1, 76) = 29.88, p < 0.0001] and Sex [*F*(1, 76) = 128.6, p < 0.0001], and a significant interaction between Treatment and Sex [*F*(1, 76) = 48.43, p < 0.0001], where EtOH preference ratio was greater in females than males for both treatments (Fig. 2C). Tukey post-hoc analysis showed a significant difference in EtOH preference ratio between saline- and CFA-treated mice for males, but not females (Fig. 2C).

**Figure 2.**
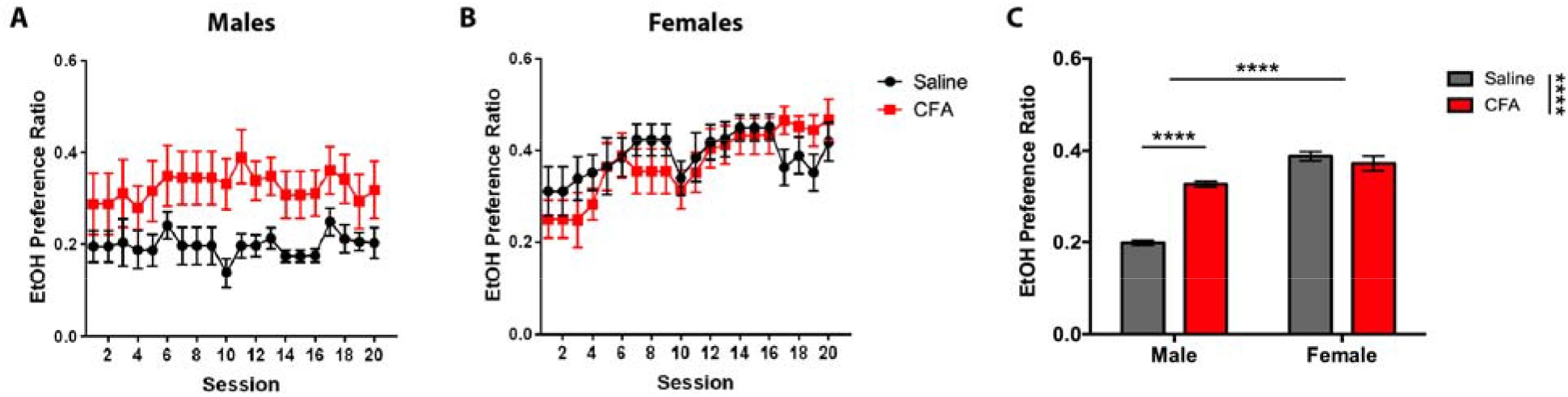
Daily alcohol preference ratio of saline- or CFA-treated C57BL/6J mice over a 3-week continuous access regimen. Pain-related alcohol preference was assessed for (A) males (n = 8) and (B) females (n = 8) in 20 consecutive sessions. (C) Averages of daily intake were compared between saline- and CFA-treated males and females. Data are mean alcohol preference ± SEM. **** p < 0.0001 difference between groups.

#### Total Fluid Intake

Despite comparable levels of total fluid intake per day, only CFA-treated females exhibited significantly greater intake than their saline-treated controls (Fig. 3). A RM-ANOVA (Treatment x Session) revealed a main effect for Session in males [*F*(19, 266) = 13.35, p < 0.0001] (Fig. 3A) and females [*F*(19, 266) = 4.514, p < 0.0001] (Fig. 3B), but no significant interaction for either sex. Sidak post-hoc analysis demonstrated no significant difference in total fluid intake between saline- and CFA-treated mice during individual drinking sessions of males or females. A twoway ANOVA (Treatment x Sex) revealed a main effect for Treatment [*F*(1, 76) = 16.38, p = 0.0001], but no main effect for Sex [*F*(1, 76) = 2.742, p = 0.1018] or significant interaction between Treatment and Sex [*F*(1, 76) = 0.2358, p = 0.6287] (Fig. 3C). Tukey post-hoc analysis showed a significant difference in total fluid intake between saline- and CFA-treated mice for females, but not males (Fig. 3C).

**Figure 3.**
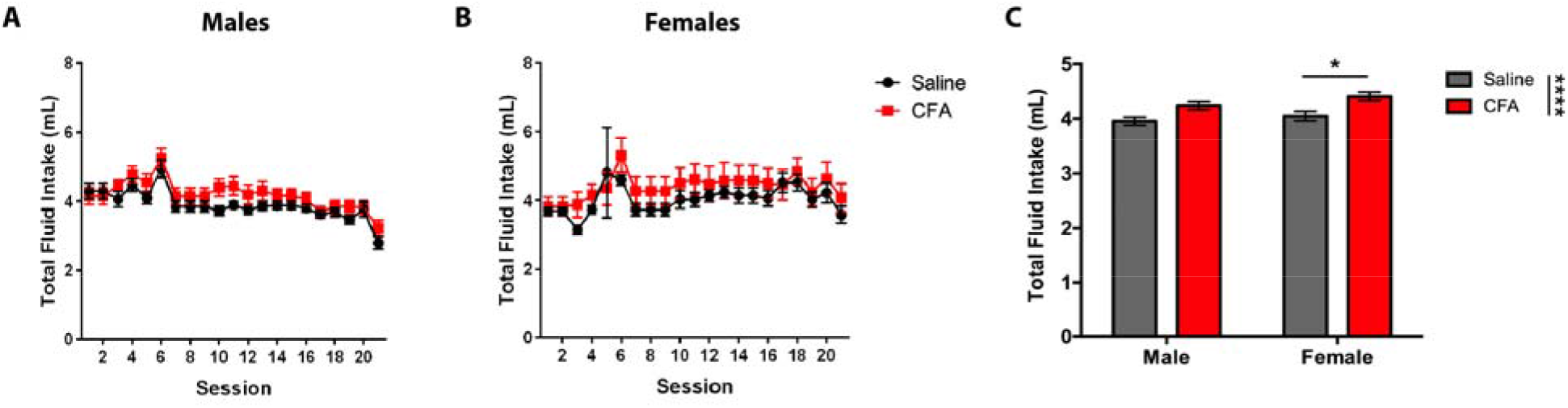
Daily alcohol and water intake (mL) of saline- or CFA-treated C57BL/6J mice over a 3-week continuous access regimen. Pain-related consumption of fluids was assessed for (A) males (n = 8) and (B) females (n = 8) in 20 consecutive sessions. (C) Averages of daily intake were compared between saline- and CFA-treated males and females. Data are mean fluid intake ± SEM. * and **** correspond to p < 0.05 and p < 0.0001 difference between groups.

### CFA-treated mice exhibit higher sensitivity to thermal nociception in both sexes

Male and female mice treated with CFA exhibit higher sensitivity to thermal nociception, as indicated by lower PWLs, when compared to saline-treated mice (Fig. 4). A RM-ANOVA (Treatment x Paw) exhibited a main effect for Treatment [*F*(1, 14) = 7.010, p = 0.0191] and Paw [*F*(1, 14) = 8.155, p = 0.0127], and a significant Treatment x Paw interaction [*F*(1, 14) = 27.46, p = 0.0001] in males (Fig. 4A), whereas a RM-ANOVA (Treatment x Paw) exhibited a main effect for Paw [*F*(1, 14) = 11.00, p = 0.0051], but not Treatment [*F*(1, 14) = 3.276, p = 0.0918], and a significant Treatment x Paw interaction [*F*(1, 14) = 10.15, p = 0.0066] in females (Fig. 4B). Sidak post-hoc analysis revealed a significant difference in PWLs between saline- and CFA-treated mice for treated paws, but not untreated paws, of both males (Fig. 4A) and females (Fig. 4B). To assess the relationship between pain and drinking, a linear regression was conducted for thermal nociceptive sensitivity of the treated paw as a predictor of EtOH intake. For males, saline-treated mice exhibited a non-significant regression equation of [*F*(1, 6) = 0.0855, p = 0.7798] with R^2^ of 0.0140 (Fig. 4C), whereas CFA-treated mice exhibited a non-significant regression equation of [*F*(1, 6) = 0.1691, p = 0.6952] with R^2^ of 0.0274 (Fig. 4C). For females, saline-treated mice exhibited a non-significant regression equation of [*F*(1, 6) = 0.0018, p = 0.9674] with R^2^ of 0.0003 (Fig. 4D), whereas CFA-treated mice exhibited a significant regression equation of [*F*(1, 6) = 8.254, p = 0.0283] with R^2^ of 0.5791 (Fig. 4D).

**Figure 4.**
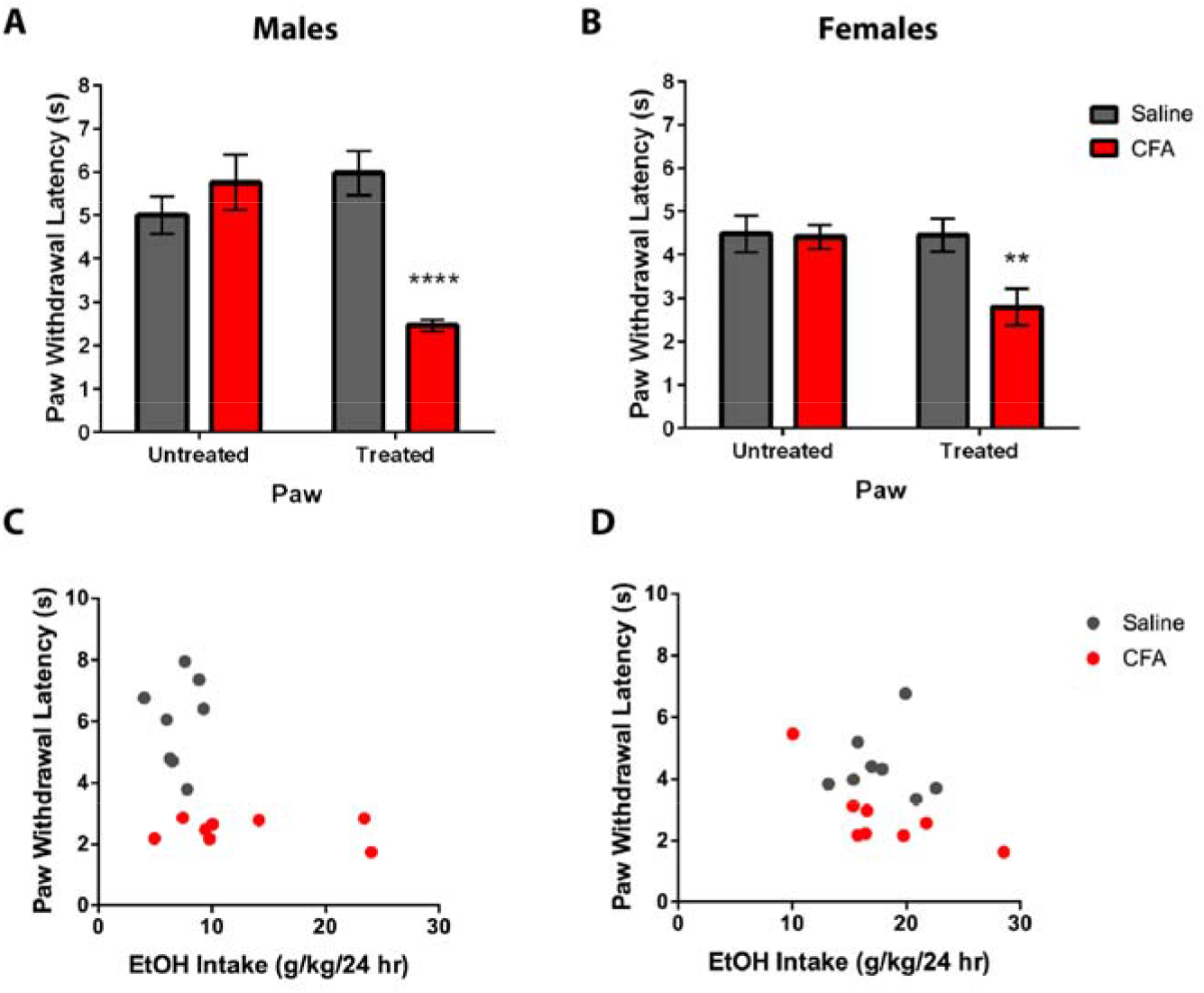
Pain sensitivity of saline- and CFA-treated C57BL/6J mice was tested 24 hrs after last alcohol exposure. Using the Hargreaves test, paw withdrawal latency (s) was measured in response to a thermal nociceptive stimulus for substance-treated and untreated paws of (A) male and (B) female mice. Pain sensitivity was correlated with average daily alcohol intake in (C) males and (D) females. Data are mean paw withdrawal latency ± SEM. ** and **** correspond to p < 0.005 and p < 0.0001 difference between groups respectively.

### CFA does not alter blood ethanol concentration and plasma corticosterone

Female mice given 2-hr access to EtOH exhibit greater alcohol and total fluid consumption than males (Fig. 5). A two-way ANOVA (Treatment x Sex) for 2-hr EtOH intake revealed a main effect for Sex [*F*(1, 28) = 83.19, p < 0.0001], but not Treatment [*F*(1, 28) = 0.1574, p = 0.6945], and no significant Treatment x Sex interaction [*F*(1, 28) = 0.6611, p = 0.4230] (Fig. 5A). Tukey post-hoc analysis showed greater 2-hr EtOH intake in female mice, but no significant difference in drinking between saline- and CFA-treated mice (Fig. 5A). A two-way ANOVA (Treatment x Sex) for 2-hr EtOH preference ratio revealed no main effect for Treatment [*F*(1, 28) = 0.2239, p = 0.6397] or Sex [*F*(1, 28) = 1.673, p = 0.2064], and a significant Treatment x Sex interaction [*F*(1, 28) = 5.599, p = 0.0251] (Fig. 5B). Tukey post-hoc analysis showed no significant difference in EtOH preference ratio between treatments or sex (Fig. 5B). A two-way ANOVA (Treatment x Sex) for 2-hr total fluid intake revealed a main effect for Sex [*F*(1, 28) = 8.726, p = 0.0063], but not Treatment [*F*(1, 28) = 0.4069, p = 0.5288], and no significant Treatment x Sex interaction [*F*(1, 28) = 1.341, p = 0.2566] (Fig. 5C). Tukey post-hoc analysis showed higher 2-hr total fluid intake in female mice, but no significant difference in drinking between saline- and CFA-treated mice (Fig. 5C).

**Figure 5.**
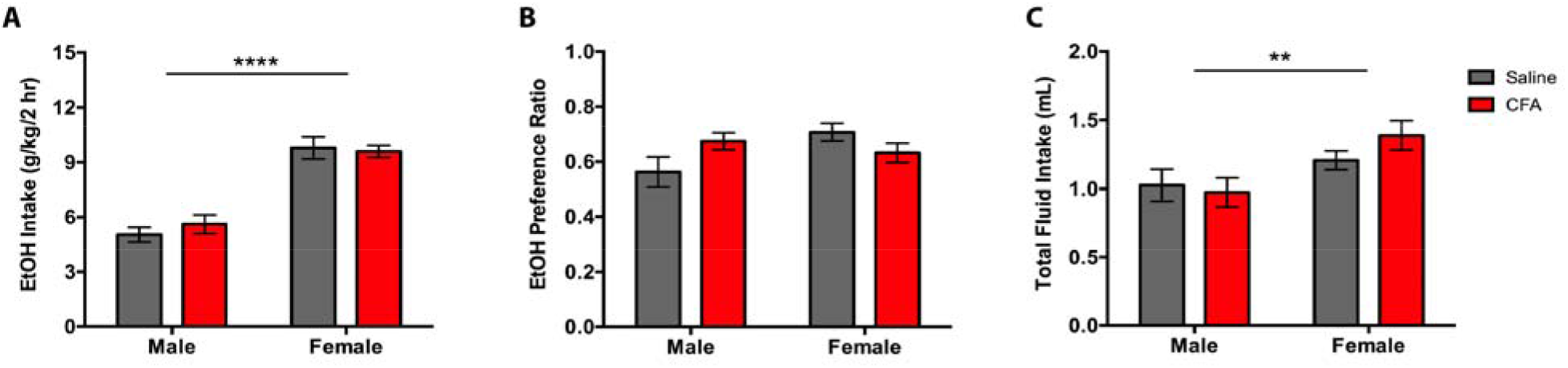
Averages of (A) alcohol intake (g/kg/2 hr), (B) preference ratio, and (C) total fluid intake (mL) in saline- or CFA-treated C57BL/6J mice during a final 2-hr drinking session. Pain-related consumption of fluids was assessed for males (n = 8) and females (n = 8) on the 21st session of a continuous access regimen. Data are mean fluid intake/preference ratio ± SEM. Data are mean fluid intake/preference ratio ± SEM. ** and **** correspond to p < 0.005 and p < 0.0001 difference between groups respectively.

Following 2-hr access to EtOH, plasma samples were collected to measure BEC and CORT levels. For BEC, a two-way ANOVA (Treatment x Sex) revealed no main effect for Treatment [*F*(1, 28) = 0.1365, p = 0.7146] and Sex [*F*(1, 28) = 0.8967, p = 0.3518], and no Treatment x Sex interaction [*F*(1, 28) = 1.890, p = 0.1801] (Fig. 6A). Tukey post-hoc analysis revealed no significant difference in BEC between treatments or sex (Fig. 6A). To assess the relationship between drinking and BEC amongst paw treatment groups, a linear regression was conducted for 2-hr EtOH intake as a predictor of BEC. For males, saline-treated mice exhibited a nonsignificant regression equation of [*F*(1, 6) = 0.0104, p = 0.9220] with R^2^ of 0.0017 (Fig. 6B), whereas CFA-treated mice exhibited a non-significant regression equation of [*F*(1, 6) = 2.334, p = 0.1774] with R^2^ of 0.2801 (Fig. 6B). For females, saline-treated mice exhibited a nonsignificant regression equation of [*F*(1, 6) = 0.5927, p = 0.4706] with R^2^ of 0.0899 (Fig. 6C), whereas CFA-treated mice exhibited a non-significant regression equation of [*F*(1, 6) = 0.6417, p = 0.4536] with R^2^ of 0.0966 (Fig. 6C).

**Figure 6.**
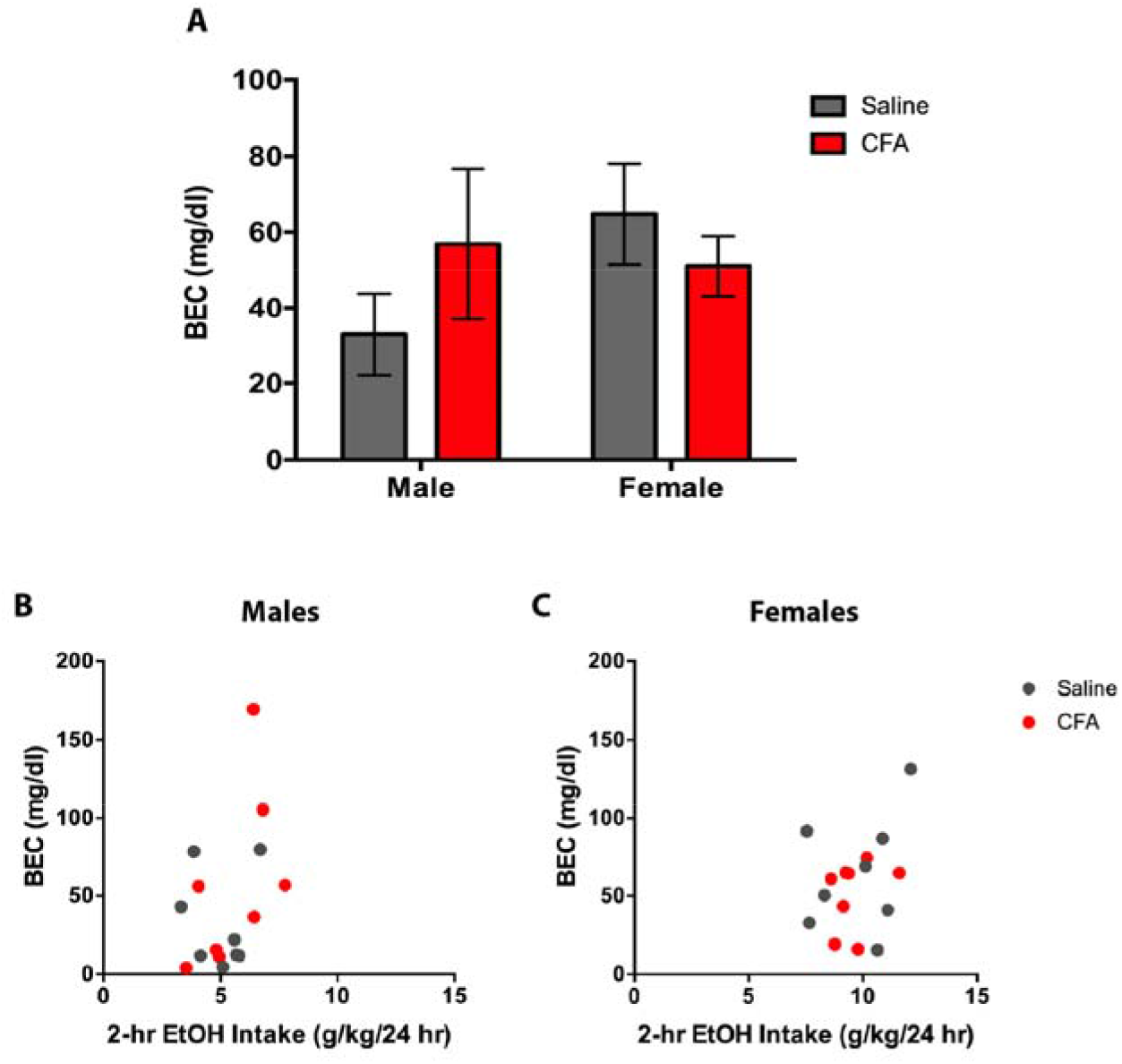
(A) Blood ethanol content (mg/dl) of saline- and CFA-treated C57BL/6J mice was measured 2-hrs into the final alcohol exposure day in male and female mice. BEC was correlated with average daily alcohol intake in (B) males and (C) females. Data are mean BEC ± SEM.

For CORT, a two-way ANOVA (Treatment x Sex) revealed a main effect for Sex [*F*(1, 28) = 7.298, p = 0.0116], but not Treatment [*F*(1, 28) = 0.1243, p = 0.7270], and no Treatment x Sex interaction [*F*(1, 28) = 0.3202, p = 0.5760] (Fig. 7A). Tukey post-hoc analysis revealed greater CORT levels in female mice, but no significant difference between saline- and CFA-treated mice (Fig. 7A). To assess the relationship between drinking and CORT amongst paw treatment groups, a linear regression was conducted for 2-hr EtOH intake as a predictor of CORT. For males, saline-treated mice exhibited a non-significant regression equation of [*F*(1, 6) = 0.3674, p = 0.5666] with R^2^ of 0.0577 (Fig. 7B), whereas CFA-treated mice exhibited a non-significant regression equation of [*F*(1, 6) = 0.0396, p = 0.8487] with R^2^ of 0.0065 (Fig. 7B). For females, saline-treated mice exhibited a non-significant regression equation of [*F*(1, 6) = 0.0016, p = 0.9691] with R^2^ of 0.0002 (Fig. 7C), whereas CFA-treated mice exhibited a non-significant regression equation of [*F*(1, 6) = 0.2297, p = 0.6487] with R^2^ of 0.0387 (Fig. 7C). Taken together, these findings suggest a weak relationship between 2-hr EtOH intake and BEC/CORT.

**Figure 7.**
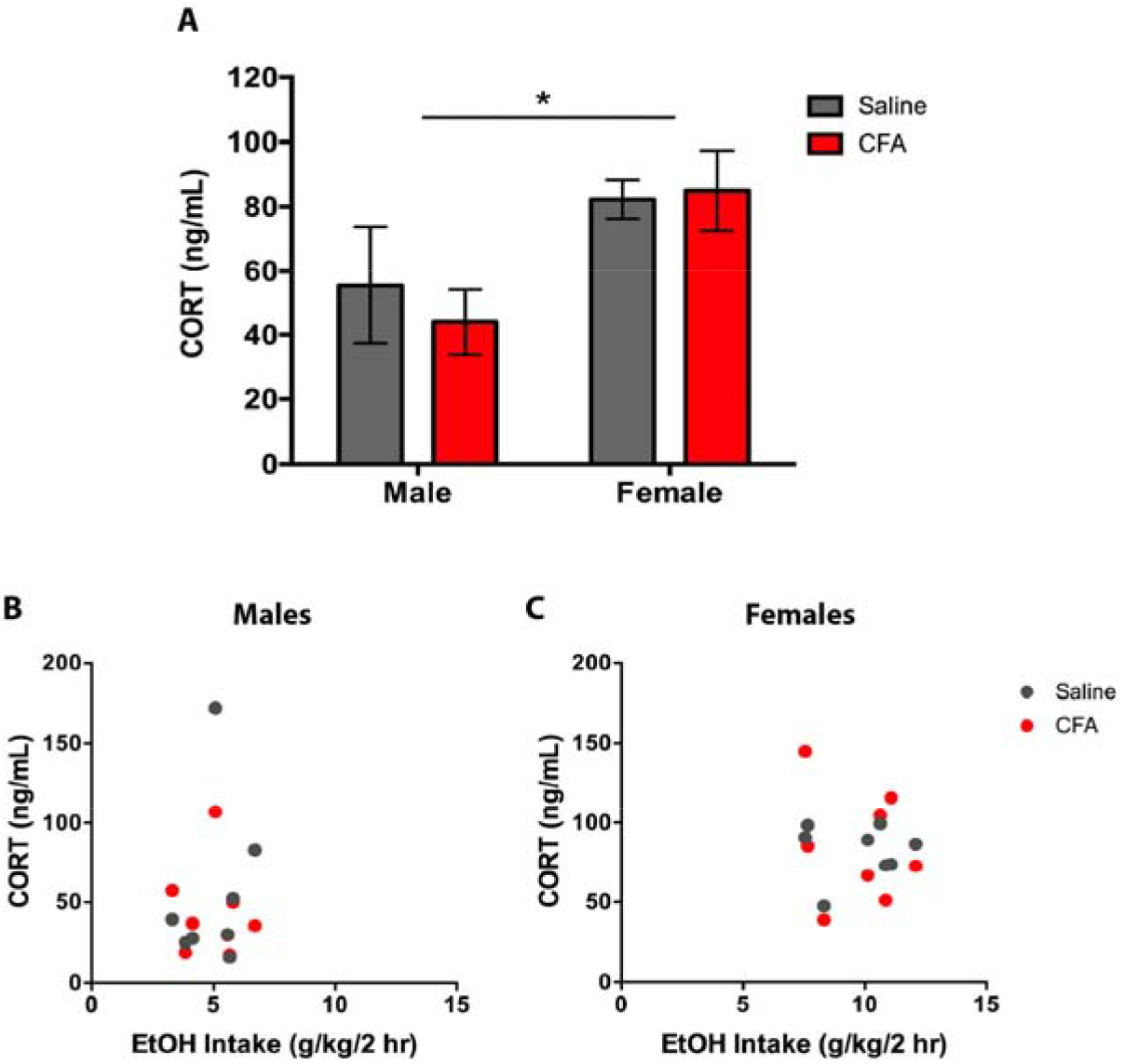
(A) Plasma corticosterone (CORT [ng/mL]) of saline- and CFA-treated C57BL/6J mice was measured 2-hrs into the final alcohol exposure day in male and female mice. CORT levels were correlated with average daily alcohol intake in (B) males and (C) females. Data are mean CORT ± SEM. * p < 0.05 difference between groups.

## Discussion

In this study, we describe sex differences in alcohol drinking following the induction of inflammatory pain. Male CFA-treated mice consumed more alcohol and exhibited higher preference for the drug than saline-treated controls, while female CFA-treated mice only showed greater total fluid intake than controls. Characteristic of C57BL/6J mice with continuous access to high concentrations of alcohol (Jury et al., 2017; Middaugh et al., 1999; Smith et al., 2015), drinking levels differed between sexes, with female mice exhibiting greater EtOH consumption, EtOH preference ratio, and total fluid intake than male mice, regardless of saline or CFA treatment. These findings suggest that chronic inflammatory pain increases alcohol drinking in males, despite higher overall drinking in females. Similar clinical observations have been made in chronic pain patients, with men more commonly self-medicating with alcohol than women (Riley and King, 2009). This pain-related alcohol consumption is greater in males throughout the lifetime, with its peak at early adulthood (Brennan et al., 2011; Riley and King, 2009; Wilsnack et al., 2009). By pairing CFA treatment and CA2BC drinking, our model was able to replicate these human patterns of voluntary drinking in injured young adult male mice, providing evidence that chronic inflammatory pain can potentiate alcohol drinking in a sex-specific manner for rodents.

Although this finding is reflective of clinical reports, it is notably inconsistent with another study on CFA-treated mice that found no effect on pain-related alcohol consumption (Smith et al., 2015). The discrepancy between findings may be due to differences in experimental design. For example, escalating concentrations of alcohol were utilized in Smith et al. (2015), while fixed concentrations were used in the present study. Mice given escalating concentrations had CFA administered while being exposed to a lower percentage of alcohol (10%), so consumption may not have produced comparable analgesia to that of a fixed high concentration of alcohol (20%). Theoretically, this would result in a higher dynamic range of alcohol consumption for the fixed concentration paradigm, which could explain why there are more pronounced effects of pain-induced drinking in the present study. Alternatively, the use of differently aged mice (i.e. 30-33 weeks old in Smith et al. [2015] versus 7-10 weeks old in the present study) could have altered alcohol consumption as well, since younger rodents drink more than fully developed adults (Holstein et al., 2011; Schramm-Sapyta et al., 2014; Vetter-O’Hagen et al., 2009). Divergences in pain severity following CFA treatment could also explain the inconsistency in findings. Male mice treated with a high volume of CFA (i.e. 50 μl in the present study) exhibited increased alcohol consumption relative to saline-treated controls, while low volume (i.e. 10 μl in Smith et al. [2015]) exposure did not alter consumption relative to pre-CFA levels. This suggests that high and low volumes of CFA treatment result in different severities of pain, which can then determine the extent of changes observed with alcohol drinking. If pain severity were to contribute to these consumption differences, a comparison of nociceptive sensitivity should reveal higher pain sensitivity in CFA-treated mice from the present study compared to those in Smith et al. (2015). The use of different behavioral assays at non-parallel time points and a lack of parallel control groups (i.e. Pre-CFA mice in Smith et al. [2015] versus saline-treated mice in the present study), however, makes it difficult to make a direct comparison. Future studies should examine how the severity of chronic inflammatory pain contributes to sex-specific patterns of drinking with more uniform measures of thermal and mechanical nociception and control groups. Determining the extent that alcohol concentration and age can alter pain-related drinking may further clarify any remaining drinking differences between the two studies.

We additionally showed that female C57BL/6J mice exhibit greater alcohol consumption than males, an observation that is common among female rodents exposed to voluntary drinking paradigms (Blanchard and Glick, 2002; Cailhol and Mormède, 2001; Chester et al., 2006; Doremus et al., 2005; Jury et al., 2017; Lancaster et al., 1996; Lê et al., 2001; Middaugh et al., 1999; Smith et al., 2015; Truxell et al., 2007). Moreover, our female drinking data matches the maximal average consumption values previously reported in C57BL/6J mice undergoing continuous access to ≥ 10% alcohol (Jury et al., 2017; Middaugh et al., 1999; Smith et al., 2015). With such high baseline alcohol consumption, it is possible that saline-treated females have already reached maximal drinking in our study, so the addition of pain would not increase drinking in females. Although this ceiling effect is possible, it is unlikely, considering that daily EtOH intake in females was around 10-12 g/kg during the first four sessions and escalated to 20-25 g/kg by the last session. If pain were able to drive alcohol drinking in females, early session EtOH intake should have increased up to two times in CFA-treated mice, but that is not the case. Therefore, sex differences in basal alcohol consumption were not likely to prevent pain-related drinking increases for females.

Differences in inflammatory response and alcohol analgesia may provide a better explanation for sex-specific patterns in drinking. CFA and related inflammatory agents cause female rodents to develop hyperalgesia more rapidly than males, with females being less prone to attenuation of inflammation by alcohol and other analgesic drugs (Alfonso-Loeches et al., 2013; Coleman and Crews, 2018; Cook and Nickerson, 2005; Pascual et al., 2017). Thus, it is possible that CFA-treated females are less susceptible to escalated alcohol consumption relative to saline-treated controls because the drug provides inadequate analgesia to promote further drinking. Although not measured in the present study, ongoing investigations with this model should assess the potential impact of sex differences in alcohol analgesia and its effects on pain-related drinking.

Three weeks after paw treatment, pain testing revealed that CFA-treated mice had higher thermal nociceptive sensitivity than saline-treated controls for both males and females. Considering the timing of the test, it is unclear whether this difference in sensitivity reflects the contributions of pain to drinking or drinking to pain. This is an important distinction, as cessation of alcohol intake following repeated drug exposure can exacerbate pain sensitivity. This phenomenon, commonly referred to as EtOH withdrawal-induced hyperalgesia (EIH), has been reported in mice (Dhir et al., 2005; Smith et al., 2016), as well as rats (Dina et al., 2000, 2007; Edwards et al., 2012; Fu et al., 2015; Gatch and Lal, 1999; Malec et al., 1987; Roltsch et al., 2017; Shumilla et al., 2005) and humans (Boissoneault et al., 2018; Dodds et al., 1945; Jochum et al., 2010; Riley and King, 2009; Wolff et al., 1942), using a variety of alcohol exposure models. In our CA2BC paradigm, there did not appear to be pathological shifts in sensitivity following three weeks of alcohol exposure. This is similar to previous continuous access paradigms in mice, which also failed to produce EIH (Smith et al., 2015). The lack of sensitivity differences following increased drinking in CFA-treated males, along with the non-significant regression of Drinking x Pain suggests that the extent of alcohol consumption had little effect on predicting pain sensitivity (and vice versa) in all groups.

The exception to this finding is CFA-treated females: despite comparable drinking patterns to saline-treated controls, CFA-treated females exhibited a significant negative correlation for Drinking x Pain, where injured females that drink more alcohol experience higher pain sensitivity. This result is reminiscent of clinical data showing that there is higher EIH severity and prevalence in females, with women being more likely to report significant recurrent pain and concurrent chronic pain conditions (Boissoneault et al., 2018).

Although our findings closely follow predicted results from the literature, not having nociceptive sensitivity measures prior to EtOH exposure makes it difficult to definitively conclude whether or not these data indicate a null effect of alcohol consumption on pain. Although a limitation in the study, it does not change our interpretation that male-specific increases in alcohol consumption follow the induction of chronic inflammatory pain. However, future experiments should be mindful of establishing baseline nociceptive sensitivity prior to the start of alcohol exposure when studying pain-related alcohol use.

Following 2 hours of CA2BC, females consumed more alcohol than males. Contrary to the increased drinking phenotype seen in CFA-treated males during 24-hr alcohol access, no treatment effects were observed. These results suggest that the terminal drinking session did not last long enough to capture peak alcohol consumption (i.e. 4+ hours into the dark cycle) (Smith et al., 2015). Although increases for alcohol drinking in the 2-hr CA2BC were specific to females, no sex effects were found for BEC. By contrast, a previous evaluation of BEC showed no sex differences for alcohol consumption after 30-minute access to 12% EtOH, but higher BECs in females (Middaugh et al., 1999). Although a common result for humans when both sexes are allowed an equal dosing of orally administered EtOH (Ammon et al., 1996; Sutker et al., 1983), such results are less common in rodents (Ho et al., 1989). This difference may be due to feeding manipulations coinciding with BEC measurements after drinking, as suggested in Middaugh et al. (1999). Although caloric mediation of alcohol consumption is well established in the literature (Rodgers et al., 1963; Rodgers, 1966; Thiele et al., 2012), sex differences in prandial metabolism and alcohol consumption are thought to be negligible, as thirst is more strongly motivating for alcohol drinking than caloric need in both sexes (Middaugh et al., 1999). Paradoxically, higher BECs in females are accompanied by enhancements in alcohol metabolism (Baraona et al., 2001; Collins et al., 1975; Mumenthaler et al., 1999), and may explain why female mice that consume a greater amount of alcohol do not produce higher BECs than males. Similar to alcohol consumption patterns in the 2-hr CA2BC paradigm, CORT was not affected by CFA, but differed by sex. Specifically, female CORT levels were found to be higher than that of males. Higher CORT release following alcohol exposure has previously been observed in female rodents, with deletion of circulating sex steroids attenuating the difference (Rivier, 1993). However, increased levels of CORT in female mice may not be the result of the drug, as alcohol drinking does not produce differences in CORT relative to consumption of water in males or females (Finn et al., 2004). Although CFA-treated males did not drink more in the 2-hr terminal drinking session, the literature would predict that an increase in drinking after 24-hr access would still not potentiate CORT release in the males, despite showing pain-related increases in alcohol consumption. This suggests that CORT is basally higher in females, and not susceptible to changes following CFA treatment in both sexes.

In conclusion, male and female mice exhibit distinct alcohol drinking behavior when undergoing pain. Compared to saline-treated controls, CFA-treated males show greater alcohol consumption, while CFA-treated females show a closer relationship between alcohol consumption and pain sensitivity. Regardless of pain state, females exhibited higher levels of total alcohol drinking and corticosterone compared to males. These findings reflect sex-specific trends in alcohol use reported by chronic pain populations and may speak to the validity of our CFA/CA2BC model in mice. Mechanistic contributions to sex differences in pain-related alcohol drinking will be required in future applications of the model, with special attention reserved for the role of stress signaling in pain and alcohol interactions.

